# Live imaging of echinoderm embryos to illuminate evo-devo

**DOI:** 10.1101/2022.08.05.503002

**Authors:** Vanessa Barone, Deirdre C. Lyons

## Abstract

Echinoderm embryos have been model systems for cell and developmental biology for over 150 years, in good part because of their optical clarity. Discoveries that shaped our understanding of fertilization, cell division and cell differentiation were only possible because of the transparency of sea urchin eggs and embryos, which allowed direct observations of intracellular structures. More recently, live imaging of sea urchin embryos, coupled with fluorescence microscopy, has proven pivotal to uncovering mechanisms of epithelial to mesenchymal transition, cell migration and gastrulation. However, live imaging has mainly been performed on sea urchin embryos, while echinoderms include numerous experimentally tractable species that present interesting variation in key aspects of morphogenesis, including differences in embryo compaction and mechanisms of blastula formation. The study of such variation would allow us not only to understand how tissues are formed in echinoderms, but also to identify which changes in cell shape, cell-matrix and cell-cell contact formation are more likely to result in evolution of new embryonic shapes.

Here we argue that adapting live imaging techniques to more echinoderm species will be fundamental to exploit such an evolutionary approach to the study of morphogenesis, as it will allow measuring differences in dynamic cellular behaviors - such as changes in cell shape and cell adhesion - between species. We briefly review existing methods for live imaging of echinoderm embryos and describe in detail how we adapted those methods to allow long-term live imaging of several species, namely the sea urchin *Lytechinus pictus* and the sea stars *Patiria miniata* and *Patiriella regularis*. We outline procedures to successfully label, mount and image early embryos for 10-16 hours, from cleavage stages to early blastula. We show that data obtained with these methods allows 3D segmentation and tracking of individual cells over time, the first step to analyze how cell shape and cell contact differ among species.

The methods presented here can be easily adopted by most cell and developmental biology laboratories and adapted to successfully image early embryos of additional species, therefore broadening our understanding of the evolution of morphogenesis.

## Introduction

Echinoderm embryos, and the sea urchin in particular, have been models for cell and developmental biology for over a century (Briggs and Wessel, 2006), leading to fundamental discoveries that shaped our understanding of fertilization (Hertwig, 1875), cell differentiation (Driesch H., 1892; Hörstadius and Horstadius, 1973; Davidson, 2006; McClay, 2011), genetic inheritance (Boveri, 1902) and cell-cycle regulation (Evans et al., 1983), to name a few. Some of these discoveries were made possible by the optical clarity of sea urchin embryos and the ease with which they can be live-imaged: these characteristics allowed, for instance, the first observations of male and female pronuclear fusion during fertilization (Hertwig, 1875) and of microtubule spindles during cell division (Hertwig, 1875). More recently, live imaging of sea urchin embryos, coupled with fluorescence microscopy, has proven pivotal to study mechanisms of epithelial to mesenchymal transition (Saunders and McClay, 2014), cell migration (Miller et al., 1995; Peterson and McClay, 2003; Campanale et al., 2014; Martik and McClay, 2015; Sepúlveda-Ramírez et al., 2018) and gastrulation (Hardin, 1988; Hardin and McClay, 1990; Kimberly and Hardin, 1998; Martik and McClay, 2017; McClay et al., 2020). Still, echinoderm embryos have so much more in store for us to discover, especially when we start shopping in the “Evolution” aisle.

Among the echinoderms, there are numerous experimentally tractable species that share a common developmental program while presenting differences in key aspects of embryonic development, e.g. asymmetry of cell divisions (Barone et al., 2022; Arnone et al., 2015; Poon et al., 2019), mode of gastrulation (Kuraishi and Osanai, 1992; Martik and McClay, 2017), presence or absence of a larval skeleton (Dan-Sohkawa and Satoh, 1978; McCauley et al., 2012; McIntyre et al., 2014; Arnone et al., 2015). While sea urchins have emerged as the main model system for echinoderms, evolutionary comparisons between sea urchin and other echinoderm species, including sea stars and sea cucumbers, are allowing us to understand how variation in the gene regulatory networks controlling cell differentiation and morphogenesis cause those differences. These types of studies have identified, for instance, genes underlying variation in asymmetric cell division (Poon et al., 2019), embryonic axes specification (Weitzel et al., 2004; Yankura et al., 2010; Peng and Wikramanayake, 2013; Hinman and Cheatle Jarvela, 2014; McCauley et al., 2015; Swartz et al., 2021), germ line formation (Fresques et al., 2014; Fresques and Wessel, 2018; Perillo et al., 2022) and skeletal cell differentiation (Hinman et al., 2003; McCauley et al., 2012; Cary et al., 2020). Such an evolutionary approach to the study of development is very powerful as it offers the opportunity not only to define the processes underlying development, but also to identify which nodes in those processes are more likely to produce a new developmental outcome, when changed. Implementing live imaging approaches for more echinoderm species would allow us to exploit the power of an evolutionary approach to aspects of morphogenesis that would otherwise be difficult to study. One example is the formation of a monolayered epithelium encircling a cavity, i.e. a blastula.

In several echinoderm species (Fig 1A), cleavage stages are followed by the formation of a hollow blastula (Fig 1B) (Newman, 1922; Dan-Sohkawa, 1976; Holland, 1981; Schroeder, 1981; Matsunaga et al., 2002; Cerra and Byrne, 2004; Arnone et al., 2015; Nesbit and Hamdoun, 2020). In all cases, the embryonic cells organize in a monolayered epithelium that separates the blastocoel from the extraembryonic fluid (Fig 1B) (Newman, 1922; Dan-Sohkawa, 1976; Holland, 1981; Schroeder, 1981; Matsunaga et al., 2002; Cerra and Byrne, 2004; Arnone et al., 2015; Nesbit and Hamdoun, 2020). However, the initial compaction of the early embryo is very variable and the blastula forms in different fashions (Newman, 1922; Dan-Sohkawa, 1976; Matsunaga et al., 2002; McCauley et al., 2012; Nesbit and Hamdoun, 2020). Sea urchin embryos, for instance, are compact until the 8-cell stage, when cell-cell contacts on the inner side of the embryo are progressively reduced and a liquid-filled blastocoel forms (Fig 1B) (McClay, 2011). In contrast, in sea star embryos, blastomeres adhere loosely to one another initially, with fluid flowing between the inside and outside of the embryo until about the 512-cell stage, when embryonic cells form large cell contacts with one another and the epithelium closes to encircle the blastocoel (Barone et al., 2022; Newman, 1922; Dan-Sohkawa, 1976; Kominami, 1983) (Fig 1B). In some cases, as in the sea star *Astropecten scoparius*, the blastomeres of the early embryo do not adhere to each other at all, but rather to the fertilization envelope (Fig 1B): the blastula is formed by blastomeres lining up along the fertilization envelope during subsequent rounds of cell division and eventually sealing the blastocoel (Matsunaga et al., 2002) (Fig 1B). Given the highly dynamic nature of blastula formation, involving changes in cell shape, cell-matrix and cell-cell adhesion, being able to perform live imaging of those different species would be an invaluable tool to identify the molecular and cellular mechanisms underlying variation in the process of forming a blastula.

**Fig. 1.**
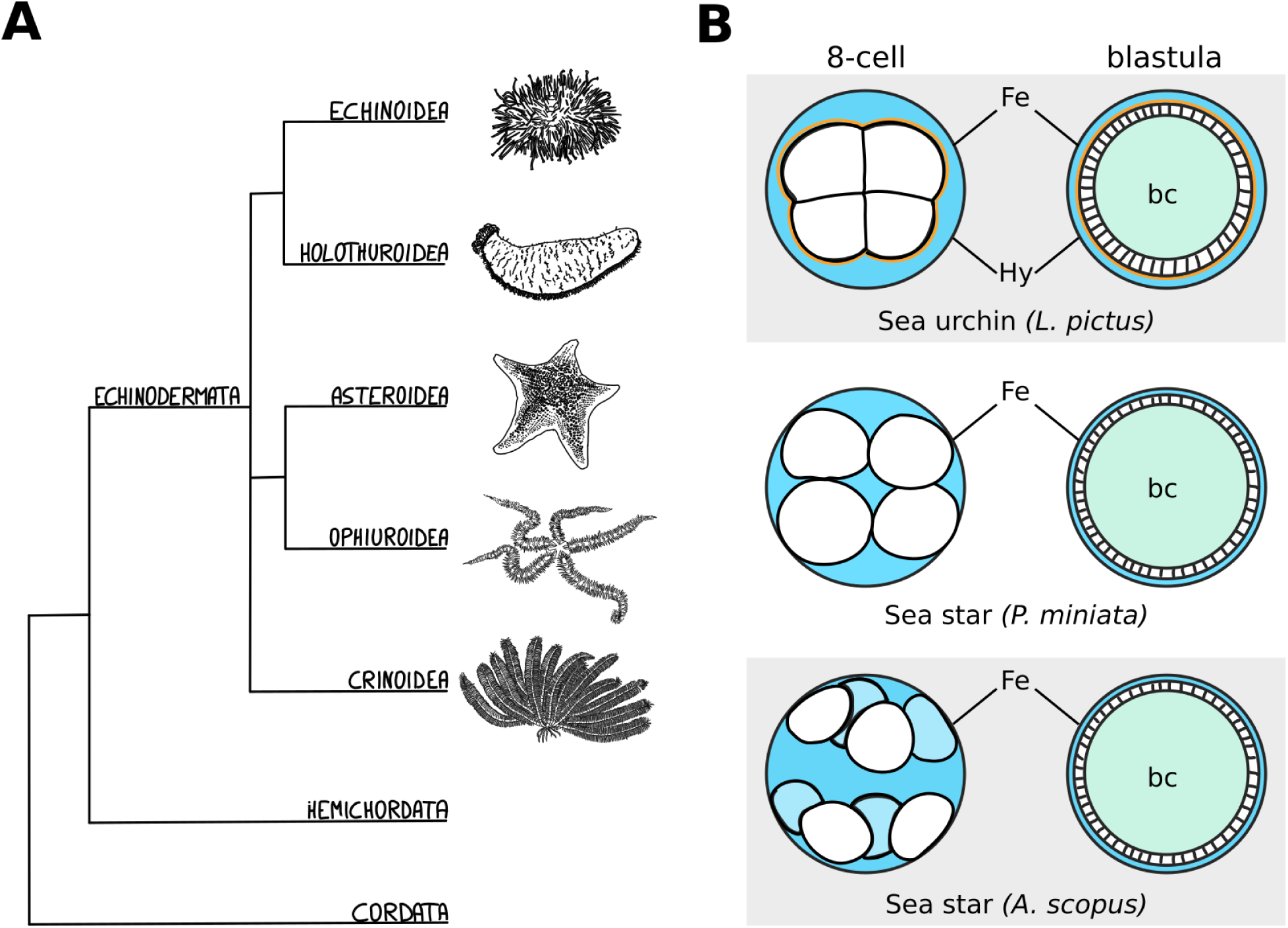
Echinoderm embryos as evo-devo models for epithelial compaction. (**A**) Schematic evolutionary tree for the major echinoderm groups. Species with transparent embryos, ideal for imaging epithelial morphogenesis, have been described among the echinoidea, holothuroidea and asteroidea. (**B**) Schematic representation of the varied mechanisms of epithelial compaction observed among echinoderms. In sea urchins the early embryo has an outer layer of extracellular matrix - the hyalin layer (Hy) - and the blastomeres are compact, adhering strongly to one another with the embryo sitting in the middle of the fertilization envelope (Fe), and extracellular fluid (blue) is excluded from the embryo. At later stages, fluid accumulates inside the embryo and forms the blastocoel (bc, teal). In sea star species, the blastomeres are less compact and the shape of the embryo is determined mainly by the size of the fertilization envelope. In some cases, as described for *A. scopus*, blastomeres do not adhere to each other but rather to the fertilization envelope itself. The sea star blastula is formed by blastomeres lining up along the fertilization envelope during subsequent rounds of cell division: eventually the embryonic cells undergo compaction and form a monolayered epithelium that seals the blastocoel.

Luckily, these echinoderm species have transparent and accessible embryos that allow for the visualization of developing epithelia at subcellular resolution (Newman, 1922; Matsunaga et al., 2002; Weitzel et al., 2004; McCauley et al., 2012; Nesbit and Hamdoun, 2020; Henson et al., 2021; Swartz et al., 2021). Optimized protocols for live imaging would permit researchers to obtain long term timelapse movies, without perturbing normal embryonic development. Here, we briefly review existing methods for live imaging of echinoderm embryos and describe in detail how we adapted those methods to allow for long-term live imaging of several species, namely the sea urchin *Lytechinus pictus* and the sea stars *Patiria miniata* and *Patiriella regularis*. We outline procedures to successfully label, mount and image early embryos for 10-16 hours, during cleavage to early blastula stages. We show that data obtained with these methods allows 3D segmentation and tracking of individual cells over time, the first step to analyze how cell shape and cell contacts differ among species.

Importantly, the methods presented can be easily adopted by most cell and developmental biology laboratories and adapted to successfully image early embryos of additional echinoderm species, and more. We recently used similar protocols to image spiral cleavage in the embryo of the snail *Crepidula atrasolea* for over 15h (Movie S1). Expanding live imaging methods to a wider number of organisms will help broaden our understanding of morphogenetic events that are, as of now, challenging to study.

### Existing methods for labeling

Understanding epithelial morphogenesis ultimately means understanding what each cell within a tissue that contributes to the morphogenetic event is doing. Direct observation of cellular dynamics – e.g. changes in cell shape and cell-cell contacts, which have greatly advanced our understanding of the physical and molecular mechanisms driving epithelial morphogenesis in model organisms (McClay, 2011; Farahani and Nelson, 2022) – will be pivotal in understanding how variation in cellular dynamics may contribute to the evolution of epithelial morphogenesis.

To visualize the dynamics of epithelial morphogenesis with cellular resolution, it is necessary to fluorescently label cellular structures so that they can be imaged live without affecting normal development. In model systems like drosophila and mouse, labeling is achieved mainly by the generation of transgenic animals where a protein localizing to the cellular structure of interest is fluorescently tagged (Garcia et al., 2011; Munjal et al., 2015; McDole et al., 2018; Dunst and Tomancak, 2019; Özgüç et al., 2022). Stable transgenic lines expressing fluorescently tagged proteins are not yet available for echinoderms, however several methods have been used to label echinoderm embryos for live imaging, with various degrees of difficulty (Strickland et al., 2004; Villoutreix et al., 2016; Campanale et al., 2019; Ortiz et al., 2019; Sepúlveda-Ramírez et al., 2019).

The easiest method is the use of vital dyes. Many vital dyes are now commercially available that stain cellular compartments, such as plasma membranes (e.g. FM or Cell Mask Orange, Invitrogen, C10045), cytoplasm (e.g, Calcein-AM, Invitrogen, 65-0853-39), acto-myosin cortex (e.g. Cell Mask Actin Tracker, Invitrogen, A57249), mitochondria (e.g. MitoView, Biotinum, 70054-T), lysosomes (e.g. LysoView, Biotinum, #70067-T), nuclei (e.g. Hoechst or Draq5, Invitrogen, 62251). Ease of use is the strong suit of vital dyes: they can simply be added to the culture medium (sea water for echinoderms) and staining is achieved in a matter of minutes (Barone et al., 2022; Campanale and Hamdoun, 2012; Sepúlveda-Ramírez et al., 2019). We have successfully used FM4 and Cell Mask dyes to label the plasma membrane and image early sea urchin embryos (Fig. 2, Movie S2) and sea star larvae (Barone et al., 2022). However, clear labeling is achieved only for relatively short periods of time (in our hands ∼3 hours at 17C), as the membrane dye will be internalized via endocytosis and soon stain the inside of the cell as much as the plasma membrane; this results in cell boundaries being detected with less contrast over time. Therefore, datasets acquired with this labeling method are useful to appreciate cellular dynamics, but usually preclude fully quantitative analysis that require, for instance, cell segmentation.

**Fig. 2.**
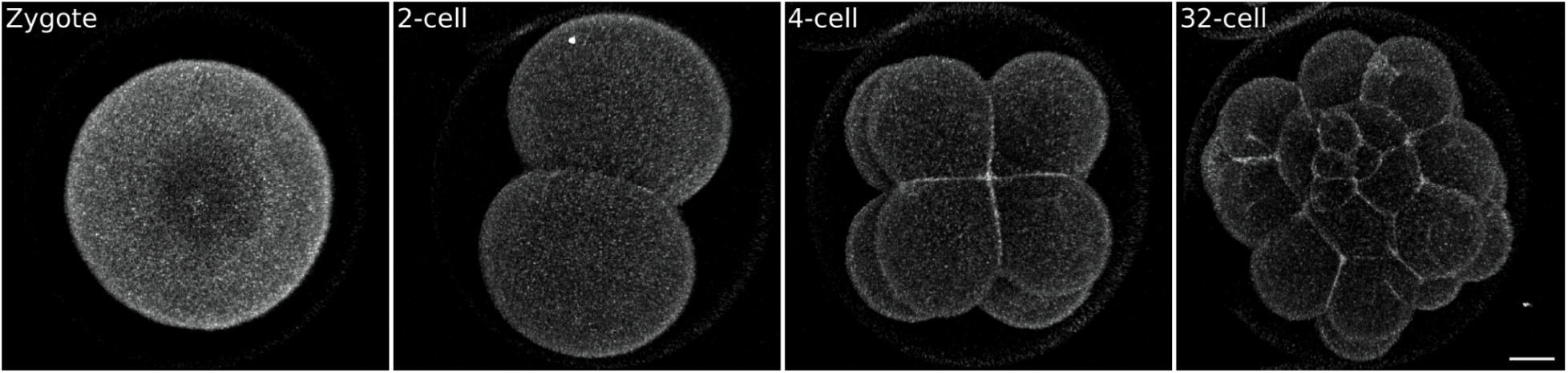
Sea urchin embryo stained with the vital plasma membrane dye FM4-64. Zygotes were mounted on a MatTek dish chamber (see Methods) in 0.5 μg/ml FM4-64 in FSW and imaged on an inverted confocal microscope.While the signal to noise ratio is sufficient to visualize embryonic cells it does not allow 3D segmentation (see Fig. 5 for comparison). Scale bar: 20 μm

In sea urchins, vital dyes can also be used to mark a specific population of cells, i.e. the micromeres (Campanale and Hamdoun, 2012; Swartz et al., 2014; Campanale et al., 2019). This is due to the fact that micromeres accumulate certain vital dyes, such as Calcein-AM, at a higher rate than other cells (Campanale and Hamdoun, 2012): the micromeres are therefore labeled more brightly than the rest of the embryo (Campanale and Hamdoun, 2012), making it possible to follow their movements (Campanale and Hamdoun, 2012; Swartz et al., 2014; Campanale et al., 2019). Another method to achieve clonal analysis is random labeling with lipophilic carbocyanine dyes, e.g. DiI (DiIC18(3)). These dyes are fluorescent lipophilic compounds that can be used to mark living cells (Barrantes, 2021). Clonal labeling of embryos can be achieved by placing DiI crystals directly in contact with the membrane of the cell to be labeled (Ruffins and Ettensohn, 1993, 1996; Henry et al., 1995) or by dissolving DiI directly in the culture medium (Ruffins and Ettensohn, 1996; Volnoukhin and Brandhorst, 2015). With the latter approach DiI will be incorporated randomly into the plasma membrane of a few cells within each embryo, effectively creating clones that can be then followed via live imaging (Ruffins and Ettensohn, 1996; Volnoukhin and Brandhorst, 2015).

Given that labeling with vital dyes does not require injection of reagents into the embryos, these methods can be readily adopted by laboratories that are not equipped with injection set-ups, including teaching labs, or by research groups interested in analyzing embryos for which injections have not yet been established. However, fully quantitative analysis allowing precise measurements of cell shapes and cell-cell contacts dynamics require labeling that remains mostly restricted to the plasma membrane, to visualize cell boundaries, and that provide high signal to noise ratios for extended periods of time. While such quality of labeling is difficult to achieve with vital dyes, it can be readily obtained driving the expression of fluorescent proteins binding to the organelle of choice (e.g the plasma membrane).

An effective method to drive the expression of a protein of interest in the whole embryo is the injection of synthetic mRNAs before the first cell division has occurred. The mRNA will be translated within the embryo resulting in ubiquitous expression of the coded protein (Gurdon et al., 1971). To label subsets of cells, mRNAs can be injected into individual embryonic cells at later stages of development (Molina et al., 2019). Injection of plasmids containing a promoter region upstream of the protein of interest is an alternative method to express fluorescent proteins. In this case the protein will be expressed only in the cells where the promoter is active: if the promoter is ubiquitous, all cells will be marked; if the promoter is specific to a cell type, only that cell type will be labeled (Barsi et al., 2014; Buckley et al., 2017; Mellott et al., 2017; Buckley and Ettensohn, 2019; Zheng et al., 2022). One important difference between these two methods is that mRNAs generally are translated in all the injected cells (Lepage and Gache, 2004), while expression from plasmids is usually mosaic (Hough-Evans et al., 1988). Therefore, injection of mRNAs coding for fluorescent proteins has been used extensively to uniformly label embryos, including echinoderm embryos (Lepage and Gache, 2004; Campanale and Hamdoun, 2012; Gökirmak et al., 2012; Campanale et al., 2014; Martik and McClay, 2017; Ortiz et al., 2019; Sepúlveda-Ramírez et al., 2019). The catalog of fluorescent proteins localizing to defined subcellular structures is vast and constantly expanding, so marking virtually any cellular organelle is possible (e.g. the plasma membrane (Gökirmak et al., 2012), the actomyosin cytoskeleton (Burkel et al., 2007), microtubules (Strickland et al., 2004; Dassow et al., 2009), centrosomes (Sepúlveda-Ramírez et al., 2019), nucleus (Villoutreix et al., 2016), etc..). Most of these markers exploit deeply conserved protein sequences and can be used in many animal species. The choice of fluorescent protein will depend mainly on the scope of the experiment: to follow individual cells over time while imaging whole tissues, we aimed at obtaining a uniform labeling of cell membranes and nuclei. In echinoderms, we have had best results with membrane bound mCitrine or GFP (lck-mCitrine and Ras-GFP) (Gökirmak et al., 2012) and tagged Histone2B (H2B-RFP, H2B-CFP) (Megason, 2009; Gökirmak et al., 2012) : these tagged proteins give uniform and clear labeling, which allows high resolution, high contrast imaging without affecting embryo development.

It is important to note that, while injection of mRNAs has been used widely for labeling early embryos, there are species in which synthetic mRNAs will not be translated. In some cases, increasing mRNA stability via the addition of a poly-adenine tail will solve the problem (McDougall et al., 2014; von Dassow et al., 2019). In fact, this step is necessary for mRNA translation in sea star embryos (von Dassow et al., 2019; Swartz et al., 2021). For species in which translation of injected mRNAs is not an option, an alternative method for labeling is the injection of previously synthesized recombinant proteins (e.g. Lifeact-EGFP (Pal et al., 2020)).

### Methods for mounting

A necessary step to achieve high resolution imaging is safely immobilizing the sample, i.e. mounting, without inflicting damage. While some movement can be corrected digitally after acquisition, best results are obtained if the sample does not move during imaging. In the case of embryos, the mounting technique needs to immobilize the embryo itself while still allowing the normal movements of cells and molecules *within* the embryo. In other words, the ideal mounting does not damage nor deform the embryo to be imaged. Especially when aiming at the study of epithelial morphogenesis, avoiding deformation of the embryo is important. Several techniques have been employed for live imaging of echinoderm embryos, foremostly applied to the sea urchin. Among these, the use of a Kiehart chamber (Kiehart, 1982), wet chambers (Martik and McClay, 2017), embedding in gels (agarose, PEG-DA) (Villoutreix et al., 2016; Burnett et al., 2018) and immobilization on an adhesive substrate such as protamine (Barone et al., 2022) have been used successfully for live imaging of the sea urchin (reviewed in (Strickland et al., 2004; Sepúlveda-Ramírez et al., 2019)). The Kiehart chamber consists of a metal scaffold in which two coverslips are kept at a defined distance from each other, forming a chamber in which embryos can be kept in place by gentle pressure (Kiehart, 1982). The chamber is then sealed with mineral oil to avoid evaporation (Kiehart, 1982). Wet chambers are adaptations of the Kiehart chamber, for short-term imaging. In this case the chamber is constructed by spacing two coverslip with a small amount of clay, loading embryos in the chamber and then sealing it with vaseline or VALAP (Strickland et al., 2004; Sepúlveda-Ramírez et al., 2019). This method also relies on slight compression of the embryos for immobilization, which is not ideal for studies of morphogenesis as it affects embryo shape.

Embedding in agarose requires placing and orienting embryos into 1% ultra-low melting agarose in filtered sea water: this solution will be liquid when kept at 20-25C and solidify into a soft gel when the temperature is lowered to 16-12C. The use of PEG-DA gels is a handy alternative that has recently been developed and tested for marine embryos (Burnett et al., 2018): in this case the polymerization of the gel is triggered not by temperature but by brief exposure to UV light. This is particularly useful for those embryos that are very susceptible to higher temperatures (even 20C), as the whole mounting procedure can be performed on a cold plate or in a temperature controlled room. In our hands, embedding methods work well for sea urchins, although it is easy to deform the embryos, especially when mounting at cleavage stages.

Coating a glass-bottom dish with protamine is a valid alternative: a 1% protamine sulfate solution is placed on the glass for a few minutes and then washed with filtered sea water. The glass-bottom dish is then filled with water and the embryos are placed onto the glass, to which they will stick. Protamine coating is best used on embryos that are still within their fertilization envelope: in this case the fertilization envelope will remain attached to the glass, effectively immobilizing the embryo within the envelope without damage or deformation. Note that If the naked (e.g. no fertilization envelope) embryo is placed directly onto the protamine, the cell-membranes in contact with it will be spreading onto the glass, which is not ideal. We have successfully used this method for long term live imaging of the sea urchin *Lytechinus pictus*.

Other species of echinoderms, however, require different methods. Sea star embryos, for instance, are softer than sea urchin embryos, so that they cannot be embedded in gels without incurring damage. Moreover, while sea urchin embryos are compact and generally sit in the middle of their fertilization envelopes, embryos of other species - e.g. *Patiria miniata, Patiriella regularis, Astropecten scoparius, Parastichopus parvimensis* - are not compact and fill the fertilization envelope from early on. In these species, even slight deformations of the fertilization envelope following adhesion onto a protamine substrate may alter the embryo morphology. Therefore, we use an adaptation of the Kiehart chamber that allows immobilizing of such embryos and it is easily accessible. This method consists in using a glass-bottom dish to create a sealed chamber (Fig. 4A-G).

**Fig. 3.**
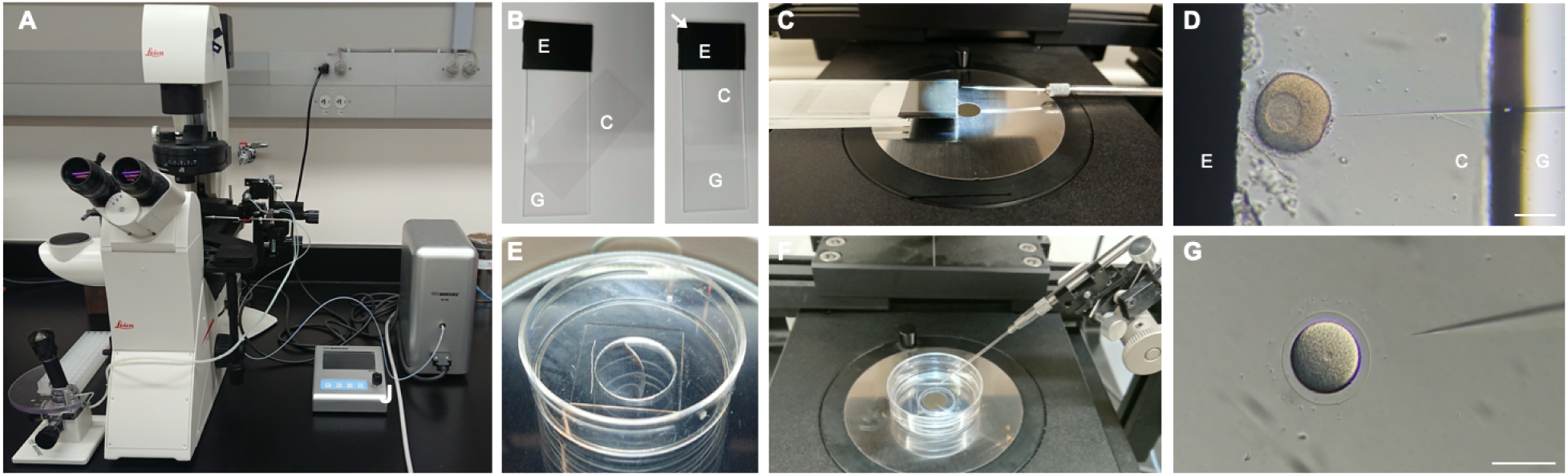
Echinoderm injection setup. (**A**) Injection set up composed of a Narishige IM-400 microinjector operated via a Narishige micromanipulator mounted on a Leica Dmi8 transmitted light inverted cell culture microscope. (**B**) Injection chamber for sea star oocytes and embryos, built by applying electrical tape (E) onto a glass slide (G) and positioning a coverslip (C) on top. (**C**) Sea star injection chamber and needle positioned for injection. (**D**) Close up view of a sea star oocyte being injected inside a chamber formed by a glass slide (G) and a coverslip (C) separated by the electrical tape (E). (**E**) MatTek glass bottom dish prepared for sea urchin injection by scraping a line in the plastic next to the glass bottom and coating the glass with 1% protamine sulfate solution. (**F**) Sea urchin injection dish and needle positioned for injection. (**G**) Close up view of a sea urchin zygote being injected inside a protamine coated glass bottom dish. Scale bars: 100 μm.

**Fig. 4.**
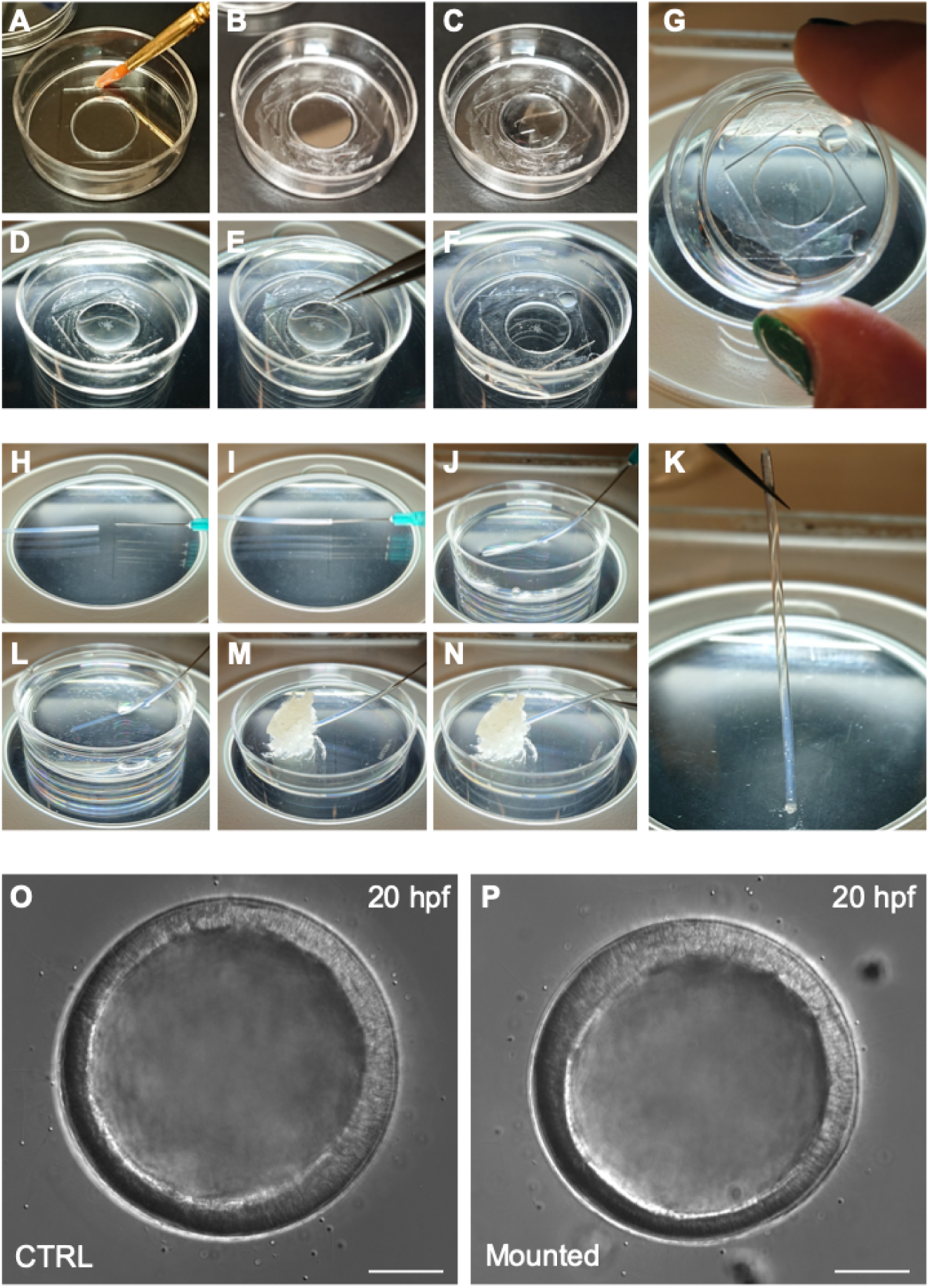
Mounting echinoderm embryos for live imaging. (**A-G**) Mounting on glass bottom dish for imaging with inverted microscopes. A MatTek dish chamber is prepared by brushing a thin layer of vaseline on the plastic portion of the MatTek dish (**A**,**B**) and adding 250 μl of PS-FSW (**C**). Embryos are then transferred to the center of the dish (**D**) and a coverslip is gently placed on top of the PS-FSW drop containing the embryos (**E**). The coverslip is pressed down slowly so that excess water is pushed out of the chamber while the embryos remain centered in the dish (**F**). At this point the chamber is sealed and capillarity keeps the embryos from moving (**G**). (**H-K**) Mounting in FEP tube for imaging on vertical light-sheet microscope. The FEP tube is connected to a syringe (**H-I**), rinsed and filled with PS-FSW (**J**). The embryos are aspirated inside the tube (**L**), which is sealed by aspirating some vaseline (**M**). The tube is then disconnected from the syringe and the second side is sealed with vaseline too (**N**). At this point the embryos are mounted within a completely sealed FEP tube and capillarity keeps them from moving (**K**). Representative images of sea star embryos cultured in a petri dish (**O**) or mounted for imaging in a MatTek chamber (**P**). Scale bar: 50 μm.

Glass-bottom dishes, such at MatTek (e.g. P35G-1.5-14-C), are regular petri dishes that have been perforated and resealed by the addition of a coverslip on the bottom of the dish, therefore allowing imaging of the specimen cultured in the dish with an inverted microscope. When mounted on such dishes, the sample is therefore located in what is practically a well within the glass-bottom dish, with diameter equivalent to the perforation and height equivalent to the thickness of the plastic. For MatTek dishes, the height is about 600 microns, which is large enough to fit most echinoderm early embryos: sealing this well with a second coverslip on the top creates a chamber in which embryos can be cultured with no damage or developmental delay for at least 20h (Fig. 4O-P). Importantly, capillarity within the chamber prevents the embryos from moving, *without the need for any mounting medium aside sea water* (Fig. 4G). We find this method is a terrific choice for imaging of particularly delicate embryos, which would be damaged or deformed by any type of mechanical constraint.

For embryos to develop properly within such a chamber, it is paramount that there is no air within the chamber and that the sealant is not toxic and does not allow any evaporation. To achieve this we use vaseline. Specifically, we brush a thin layer of vaseline on the plastic bottom of the dish, around the bottom coverslip. We then add filtered sea water to cover the bottom coverslip and place the embryos in the middle of the well. Finally, we place a second coverslip on top of the well and press it down until all excess water is pushed out of the chamber and the vaseline is uniformly adhering to all sides of the top coverslip. At this point the embryos are immobilized by means of capillarity (Fig. 4G). More water should be added in the dish to prevent any evaporation and to help maintain the desired temperature in the chamber.

Interestingly, the same principle can be used to mount live echinoderm embryos into transparent plastic tubes, FEP tubes, which allows imaging on vertical light-sheet microscopes, such as the Zeiss Z1 (Weber et al., 2014). In this case, the FEP tube is first mounted on a syringe needle, which is used to aspirate in the tube first filtered sea water containing the embryos and then a bit of vaseline to seal the tube (Fig. 4H-M). The tube is then gently detached from the syringe and sealed with vaseline on the other side (Fig. 4N). It is important to note that live echinoderm embryos mounted in this fashion will be kept still by capillarity, so that they will be held in position even when the tube is arranged vertically (Fig. 4K). This does not work for fixed embryos or for heavier embryos (such as yolk rich embryos of direct developing species) that will fall to the base when the FEP tube is held vertically. In this case, vaseline should be avoided, as it is autofluorescent, and ultra-low melting agarose can be used to seal the tube. When using ultra-low melting agarose it is important to make sure that embryos do not come in contact with the warm agarose while sealing the FEP tube, as the temperature shock will impair development.

One caveat of this method is that the chamber is completely sealed and therefore does not allow for gas exchange, which could result in hypoxia. In our hands, this is not a problem for sea star embryos, as long as the number of embryos mounted in the chamber is small (less than 20 in a MatTek dish, up to 4 in a FEP tube). It is possible that hypoxia becomes a concern for species that have larger embryos or higher metabolic rates, and so viability within the chamber should be tested before imaging a new species.

### Existing options for temperature control

Temperature control is paramount not only to ensure proper development of embryos, but also to obtain reproducible results when performing live imaging. The optimal temperature for echinoderm embryos depends on the species, with some needing precise temperature control (Foltz et al., 2004; Hodin et al., 2019a) and others tolerating quite a wide range of temperatures (*Lytechinus pictus* can be raised between 12C and 23C, (Nesbit and Hamdoun, 2020)). The three species we have used for long term live imaging tolerate temperatures up to 20C, with *P. miniata* being most sensitive and preferring a temperature of 16C. In all cases, therefore, temperature control devices that allow cooling below room temperature are needed to successfully image those embryos. This can be achieved with temperature controlled stages, installed on the microscope to be used and connected with a Peltier temperature exchange device. In this case the sample is placed onto a cooled surface and therefore kept at temperature. Several such stages are commercially available - eg. from OKO lab, CherryTemp, PeCon - and should be selected based on the specific microscope to be used. We have adopted a light-weight solution from OKO lab (H101-LG), which consists of a small chamber to be mounted directly on the stage and can be used also for weight-sensitive setups, such as piezo stages. It is important to note that i) these types of stages do not allow very uniform temperature control, as heat exchange between the stage and warmer air in the room creates temperature gradients within the sample and ii) there are considerable differences between the temperature of the stage and inside the dish. Therefore, it is important to check that the temperature settings allow maintaining the embryos at the desired temperature: for instance, we have found that setting our OKO stage at a temperature of 12C results in 16C-17C inside the dish.

Devices for temperature exchange and temperature controlled stages, however, can be rather expensive, and not available to labs that may want to get started with live imaging of echinoderm embryos. An alternative is ambient temperature control: either the room in which the microscope is located or a chamber built around the microscope can be cooled to the required temperature. In some cases, a bit of creative problem solving goes a long way. For instance, we have successfully performed live imaging of *Patiriella regularis* embryos on a confocal microscope equipped for live-imaging of mammalian cells, i.e. with a temperature control chamber to be set at 37C. We repurposed the chamber by connecting it to a portable AC system and testing optimal settings to maintain 18-20C temperature inside our petri dish.

### Imaging

Embryos labeled and mounted with the methods described above can be imaged on several microscopes, including epifluorescence, stereoscope, confocal and light sheet. The optimal microscope to be used will depend on the sample, i.e. which species of echinoderm is to be imaged, and on the level of spatiotemporal resolution to be achieved. We have imaged embryos of all three species, *Lytechinus pictus, Patiria miniata*, and *Patiriella regularis*, on laser scanning confocal microscopes with very good results (Fig. 5, Movies S3-5). Confocal microscopy is definitely an excellent choice, as it allows imaging of several embryos at once, over time, generating datasets that can be comfortably processed on a good desktop computer (we use a custom built Image analysis PC equipped with AMD Ryzen 7 3800X 3.9 GHz 8-Core, 128 GB RAM, Geforce RTX 2060 Super 8 GB, for a total cost of $2000 .ca).

**Fig. 5.**
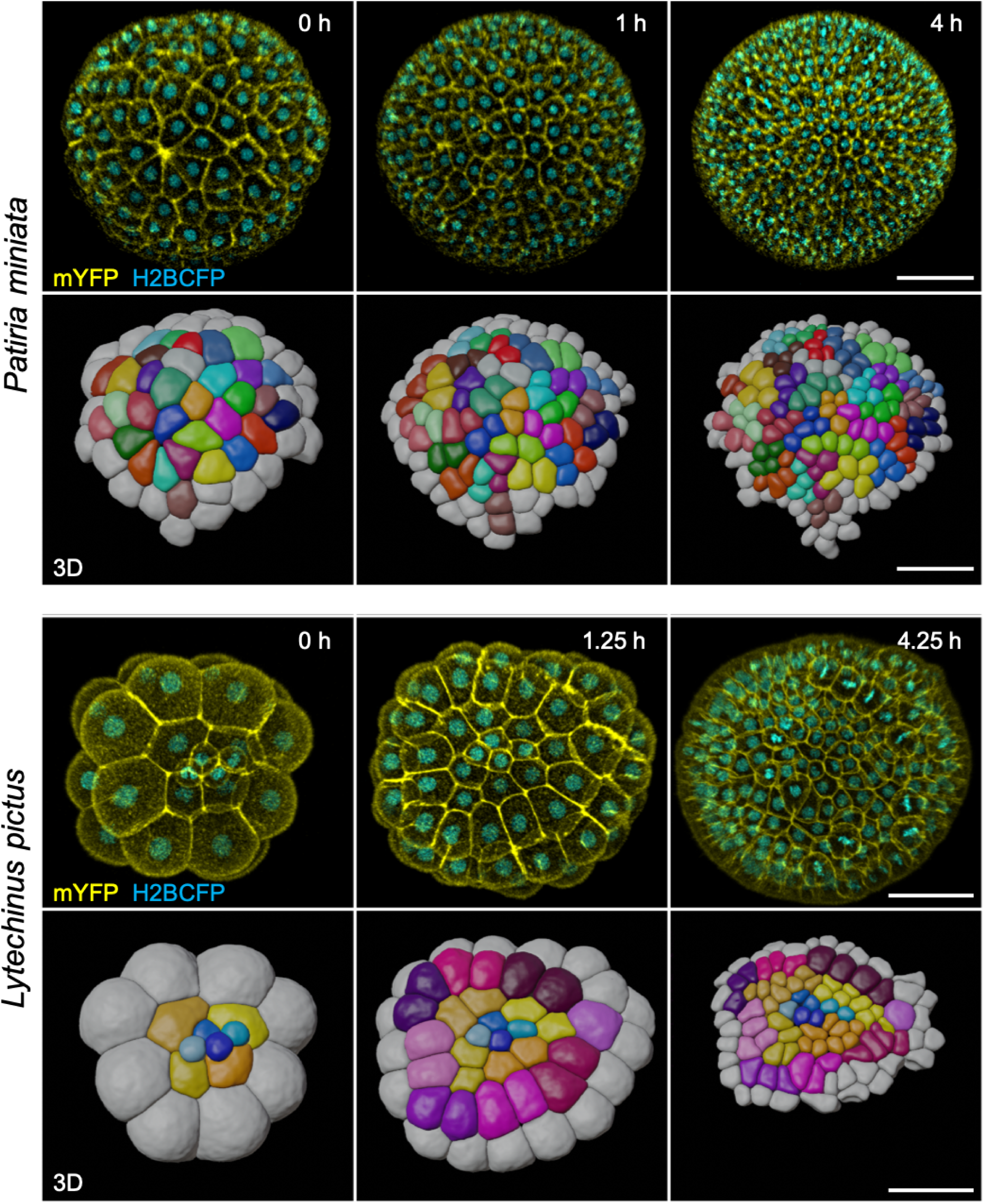
Representative stills from live imaging timelapses of sea star (*Patiria miniata)* and sea urchin (*Lytechinus pictus*) embryos. Embryos were injected, mounted and imaged on an inverted confocal microscope (see Methods). The high image quality allowed cells to be fully segmented in three dimensions and tracked using the Fiji plug-in Limeseg (shown in the 3D rows for both sea star and sea urchin). Color coding in the 3D reconstruction shows clonal relationships between the cells. Scale bars: 50 μm.

Confocal microscopy, however, does not allow imaging of the whole embryo, as the signal to noise ratio necessary for efficient segmentation and tracking of cells in 3D can be achieved for about half of a sea star embryo (embryo diameter of 150-200 microns .ca) and a little more than that for sea urchin embryos (embryo diameter of 100 microns .ca).

The use of multi-view light-sheet microscopy is recommended to obtain accurate 3D data of entire embryos (Dunst and Tomancak, 2019). It is important to note that multi-view light-sheet microscopy also requires data analysis pipelines and computing hardware that can handle very large datasets. Therefore, the imaging method of choice will depend on the type of information to be collected and the resources available for handling imaging datasets.

Independently of the microscope to be used, tuning of imaging settings will be necessary to obtain high quality images of developing embryos. Perhaps the most important parameter is the laser power used for excitation: it should be minimal. Second only to temperature, phototoxicity is the primary cause of abnormal development in embryos that are being live imaged (Sepúlveda-Ramírez et al., 2019). It is highly recommended to keep laser power as low as possible, especially when exciting with two or more laser lines at the same time, to image several fluorophores. Other parameters, e.g. zed and time resolution, will depend on the objective, microscope and information to be extracted from the datasets.

In the next section, we describe in detail the methods we used to image both sea star and sea urchin embryos from cleavage stages to hatching, providing step by step protocols for injection, mounting and live imaging. We then showcase the results obtained for *L. pictus* (Movie S3), *miniata* (Movie S4), and *P. regularis* (Movie S5), including an example of 3D cell segmentation and tracking for *L. pictus* and *P. miniata* (Fig. 5).

## Methods

### Live imaging of sea star embryos (*P. miniata* and *P. regularis*)

Adult *Patiria miniata* were purchased from Monterey Abalone Company (Monterey, CA) or South Coast Bio-Marine LLC (San Pedro, CA) and held in free flowing seawater aquaria at a temperature of 12-16C. Adult *Patiriella regularis* were collected off the coast of Tasmania (Australia) and held in aquaria at a temperature of 20C. Sea star gametes were obtained as previously described (Hodin et al., 2019b). Briefly, ovaries and spermogonia were dissected via a small incision on the ventral side of adults. Sperm was stored undiluted at 4C while ovaries were fragmented to release oocytes in FSW. Maturation of released oocytes was induced by incubating for 1h at 16C (for *P. miniata*) or 20C (for *P. regularis*) in 3 μM 1-Methyladenine (Fisher Scientific, 5142-22-3). All embryos were raised in 0.22 μm filtered sea water (FSW) with the addition of 0.6 μg/ml Penicillin G sodium salt (Millipore Sigma, P3032) and 2 μg/ml Streptomycin sulfate salt (Millipore Sigma, S1277).

mRNAs were synthesized with the mMessage mMachine SP6 Transcription Kit (Invitrogen, AM1340) and additionally polyadenylated with a PolyA Kit (Invitrogen, AM1350). *P. miniata* and *P. regularis* immature oocytes were injected with mRNAs to label membranes (lck-mCitrine or HRAS-GFP, 400 ng/μl) and nuclei (H2B-RFP, 400 ng/μl; H2A-mCherry, 400 ng/μl). Injected oocytes were incubated at 16C (*P. miniata)* or 20C (*P. regularis*) overnight, activated and fertilized.

Labeled embryos were mounted on a glass bottom dish. No medium except PS-FSW was used to immobilize the embryos: the glass bottom part of the dish was covered with a coverslip and sealed with vaseline. This creates a 600 μm deep chamber in which capillarity prevents the embryos from moving, until they develop cilia. Additional FSW was added in the dish, to help with temperature control. Embryos were incubated until the 2 cell stage and then imaged on an inverted Leica Sp8 confocal microscope (20X objective, NA 0.7, 16C controlled temperature for *Patiria miniata*) or Zeiss LSM 800 confocal microscope (20X Objective, NA 0.8, 20C controlled temperature for *Patiriella regularis*). Datasets were 3D rendered using Imaris 6.4 (Bitplane), and segmentation and tracking was achieved using the Fiji plugin Limeseg (Machado et al., 2019).

A step by step protocol is provided below.

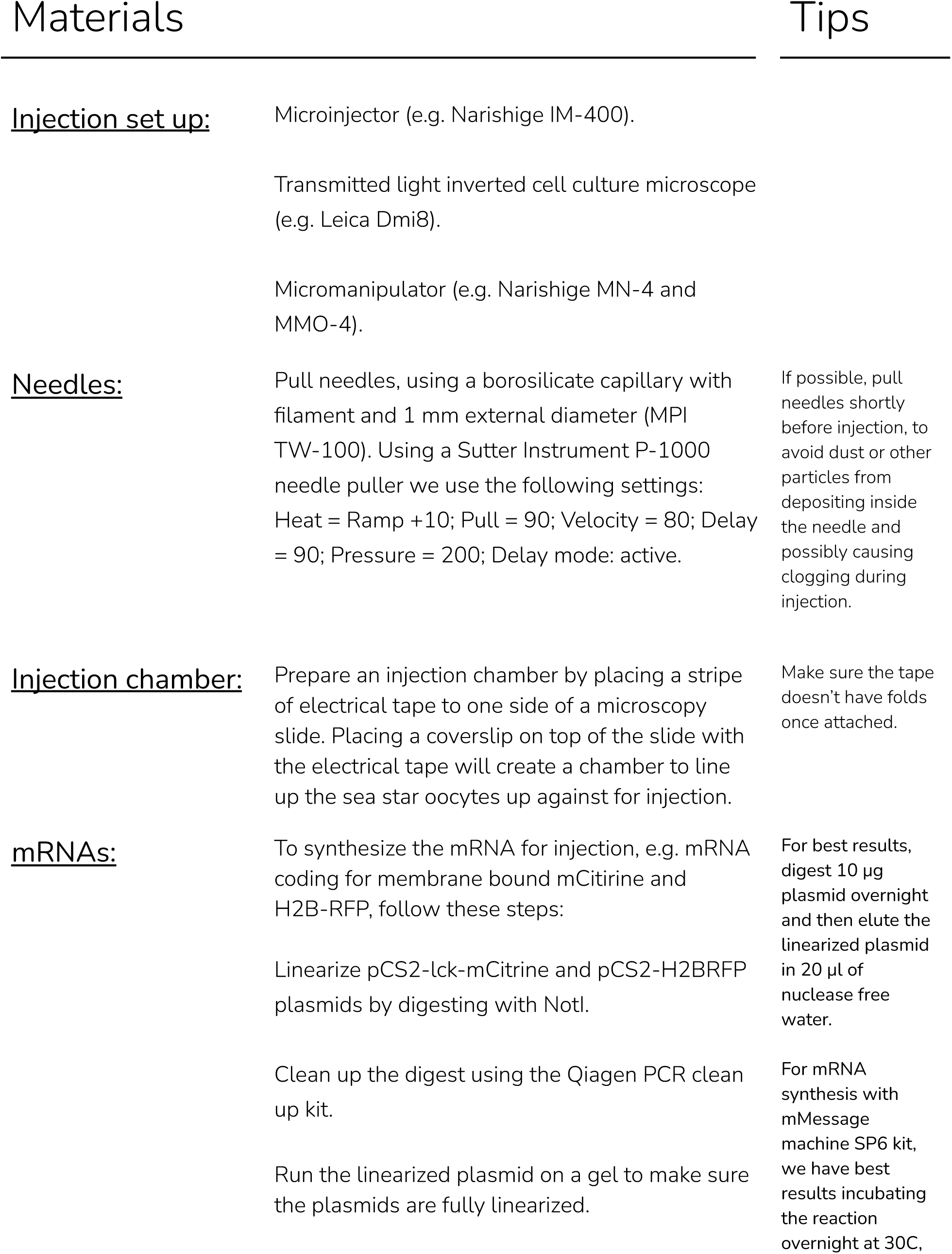
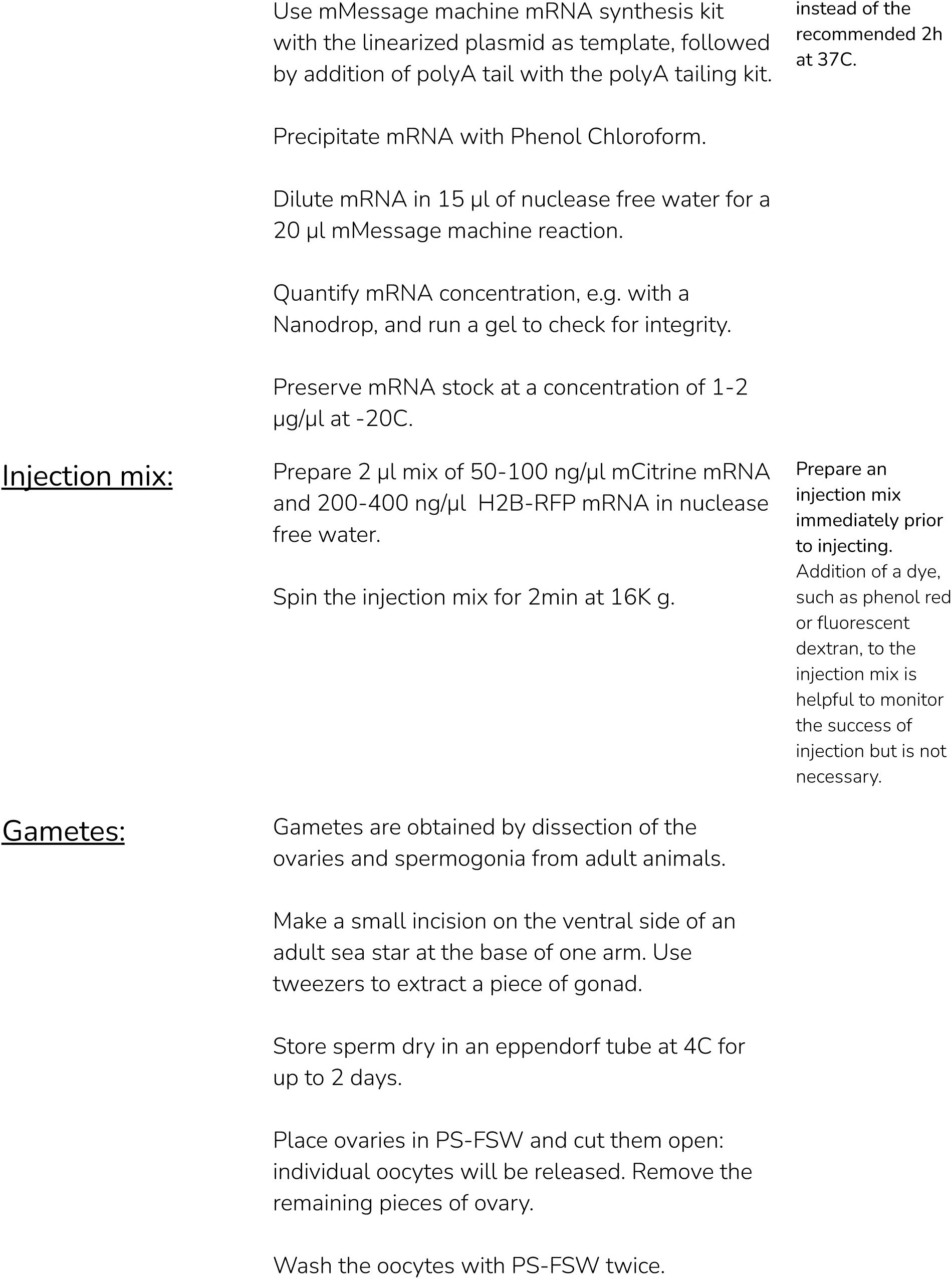
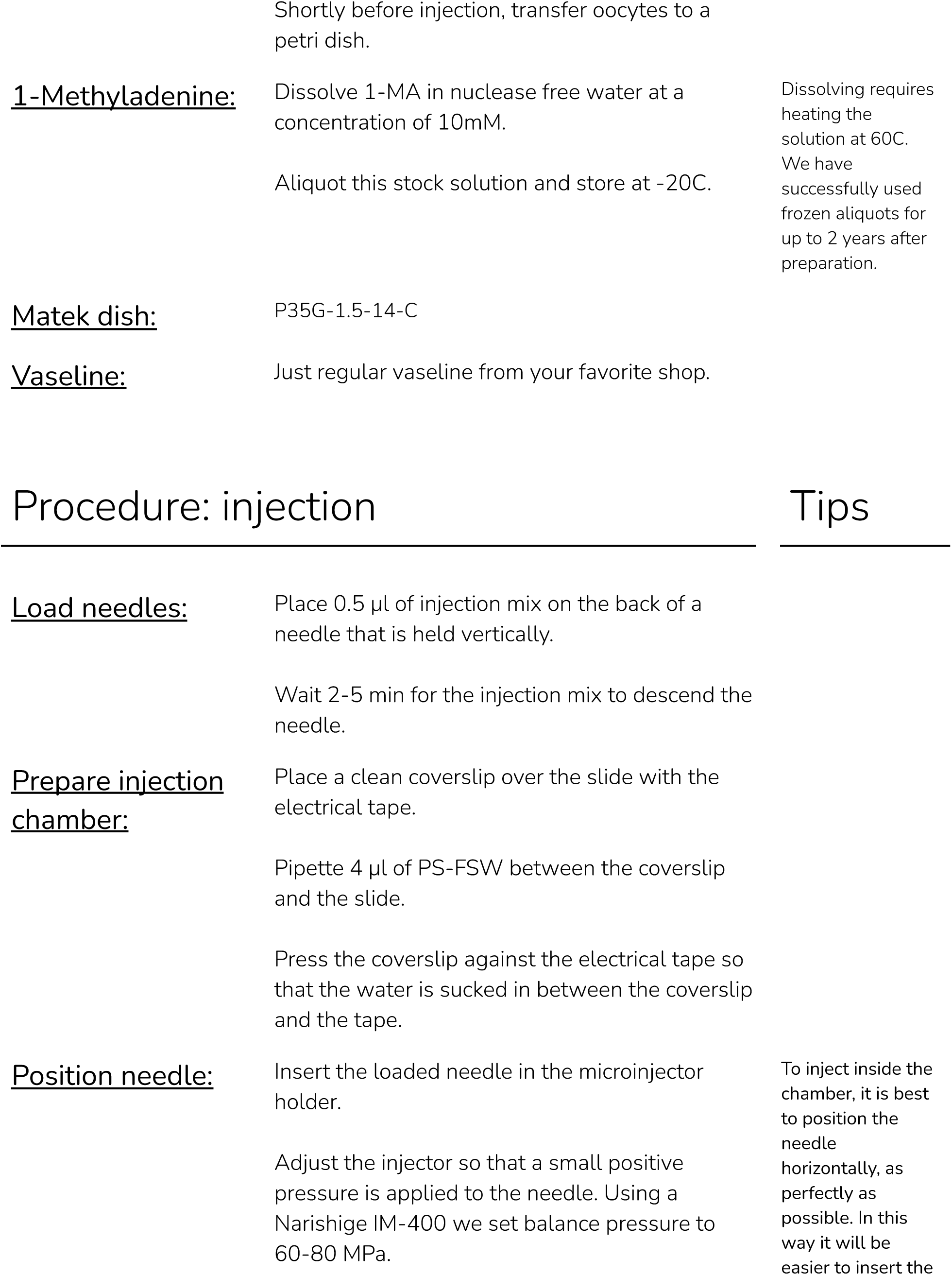
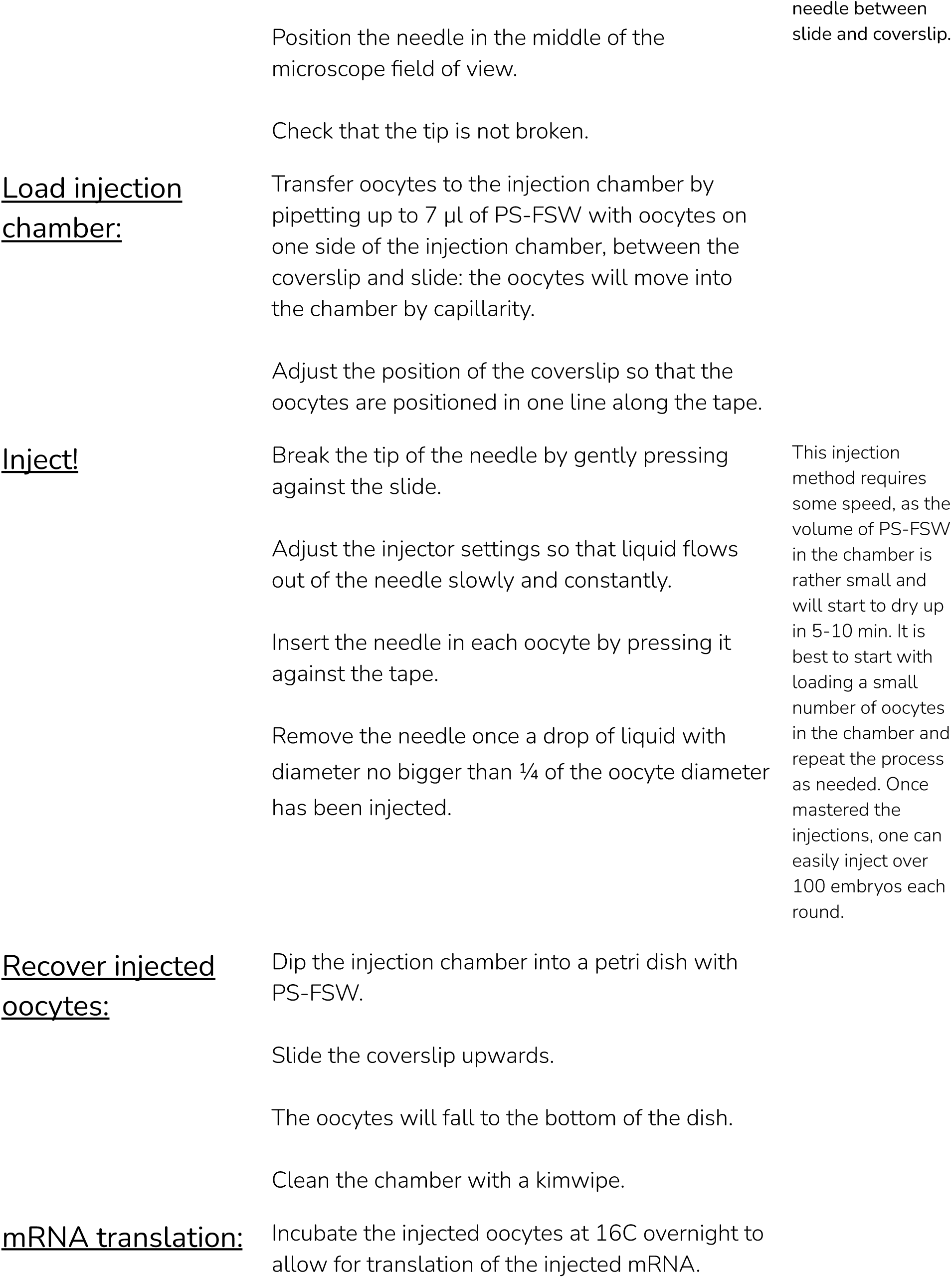
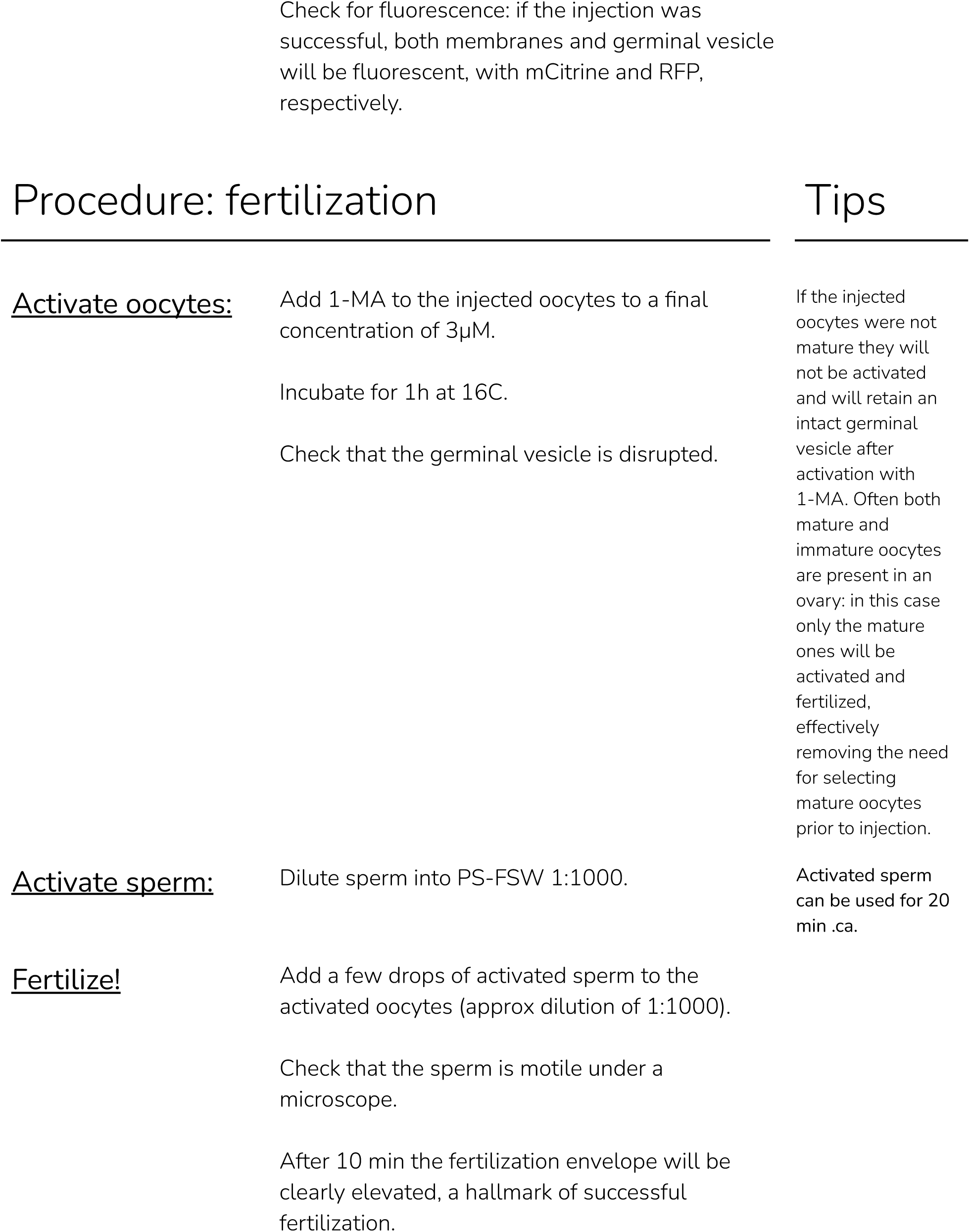
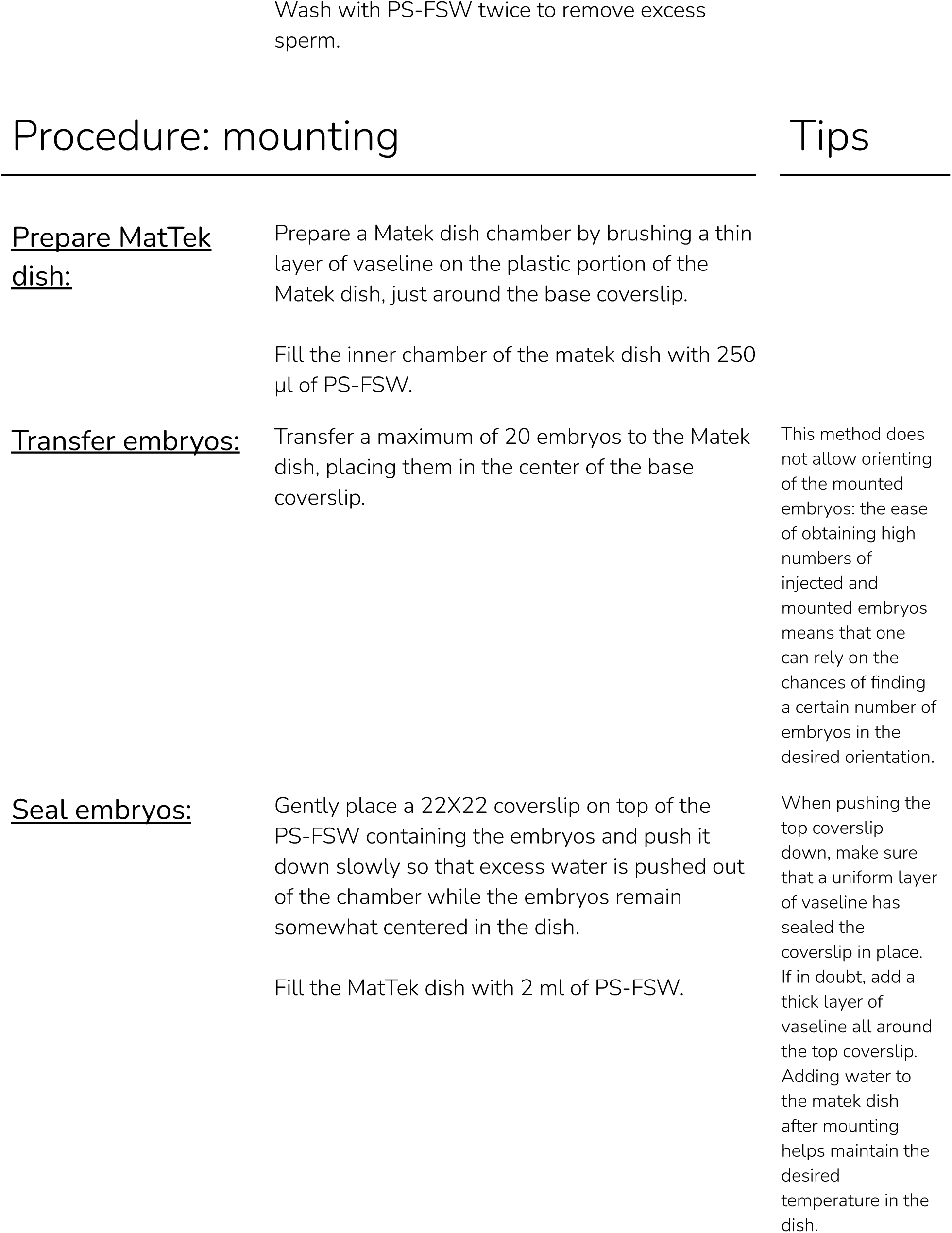
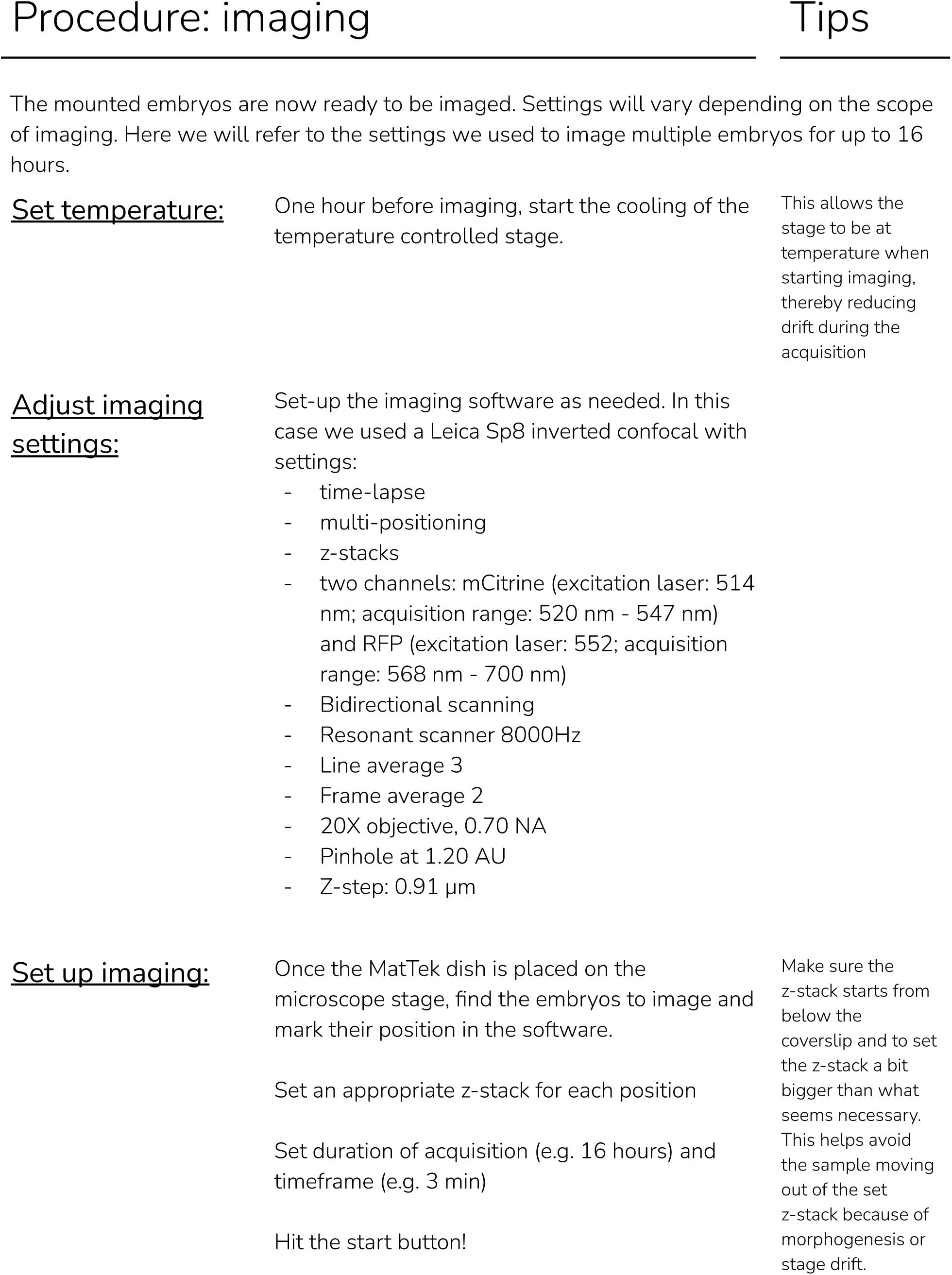

### Live imaging of sea urchin embryos

Adult *Lytechinus pictus* were collected at La Jolla, CA, and held in free flowing seawater aquaria at a temperature of 16C. Spawning was induced by injection of 0.5M KCl, as previously described (Nesbit and Hamdoun, 2020).

mRNAs were synthesized with the mMessage mMachine SP6 Transcription Kit (Invitrogen, AM1340). *Lytechinus pictus* were injected at the 1 cell stage with a mix of mRNAs coding for membrane bound mCitrine and Histone-2B-RFP (lck-mCitrine, 50 ng/μl; H2BRFP 400 ng/μl) on a glass bottom dish (MatTek, P35G-1.5-14-C) coated with protamine, incubated at 16C until the 2-cell stage and then imaged on an inverted Leica Sp8 confocal microscope (20X objective, NA 0.7, 16C controlled temperature) until the blastula stage. Datasets were 3D rendered using Imaris 6.4 (Bitplane), segmentation and tracking was performed using the Fiji plugin Limeseg (Machado et al., 2019).

A step by step protocol is provided below.

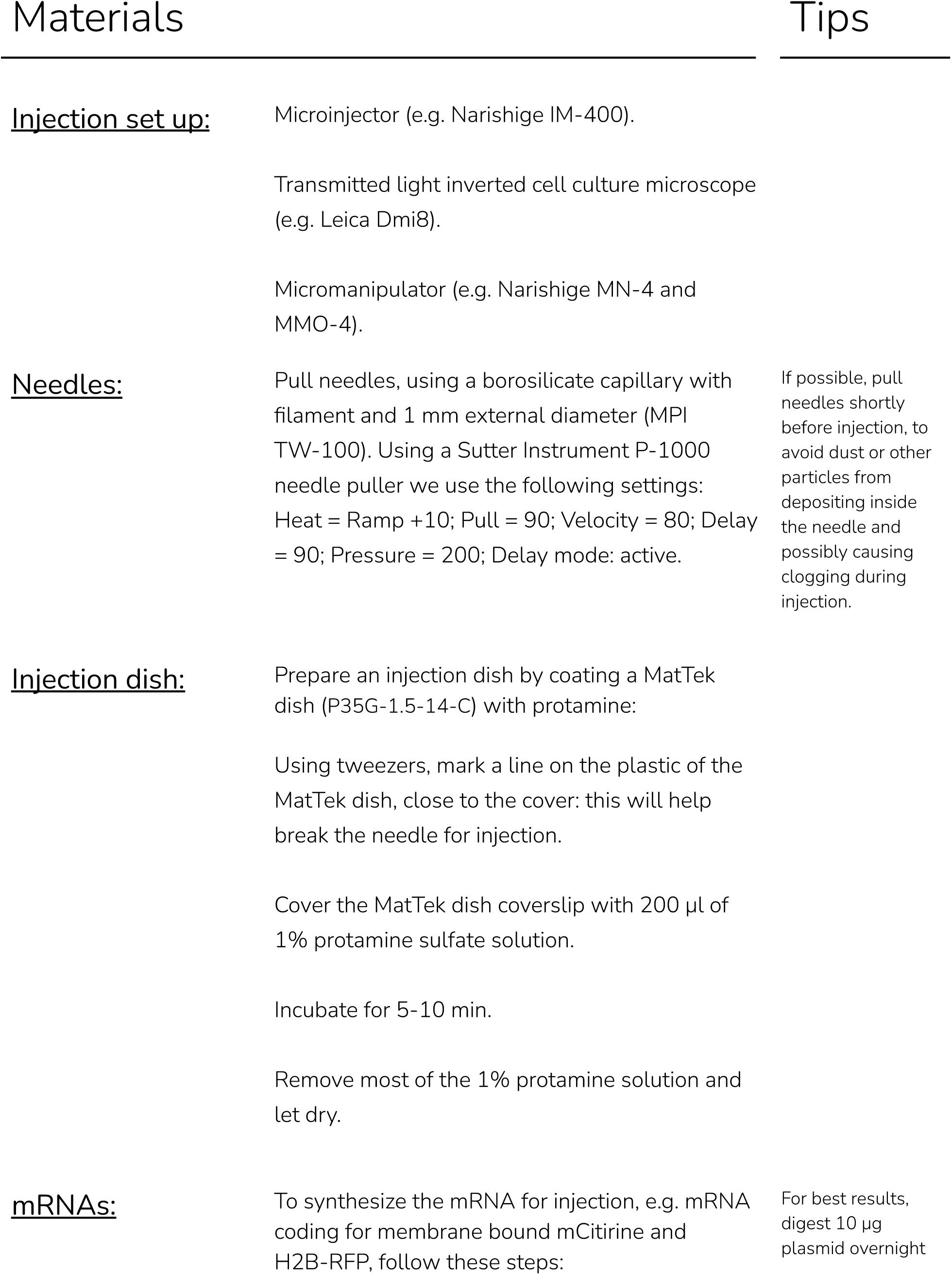
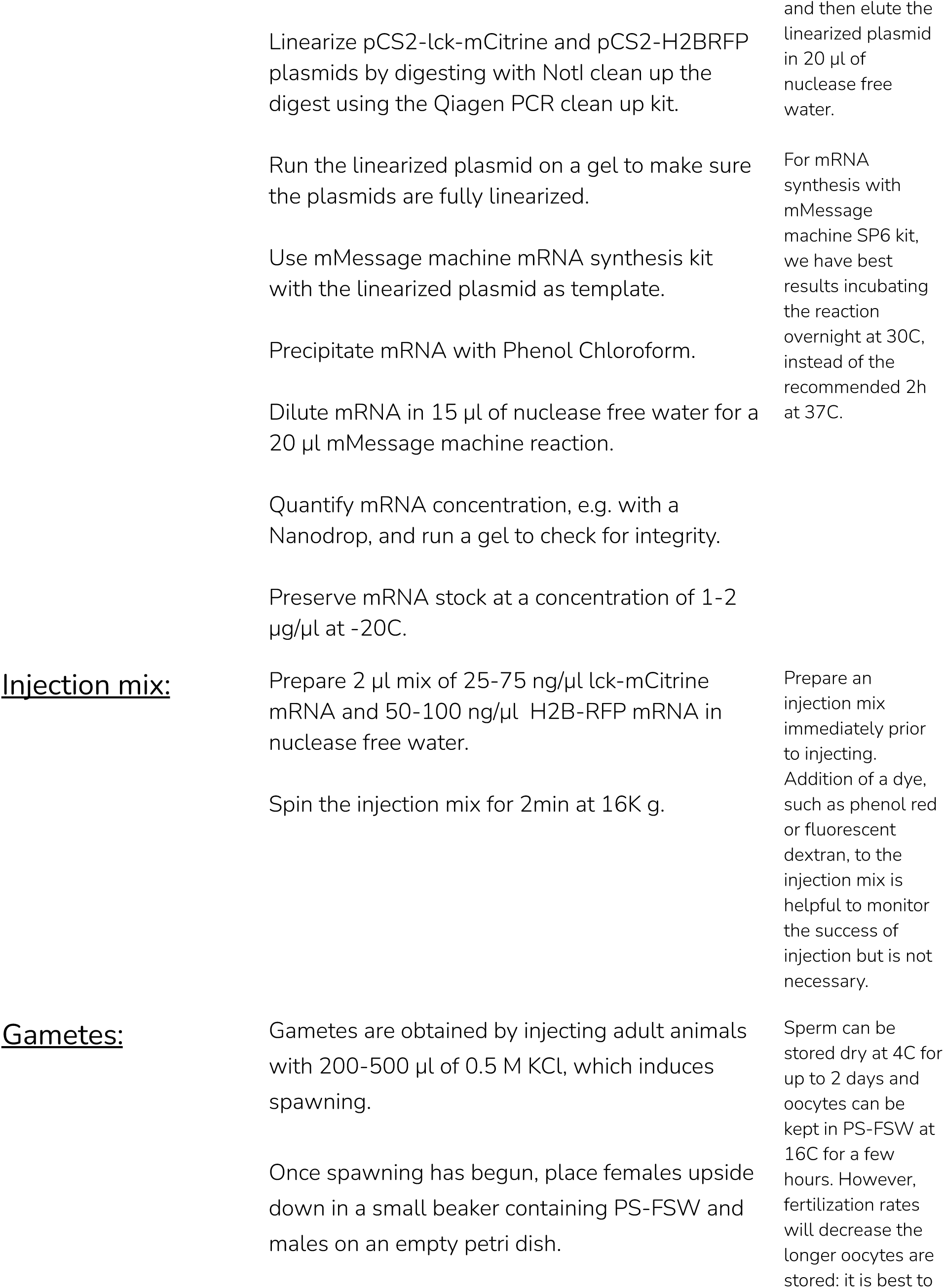
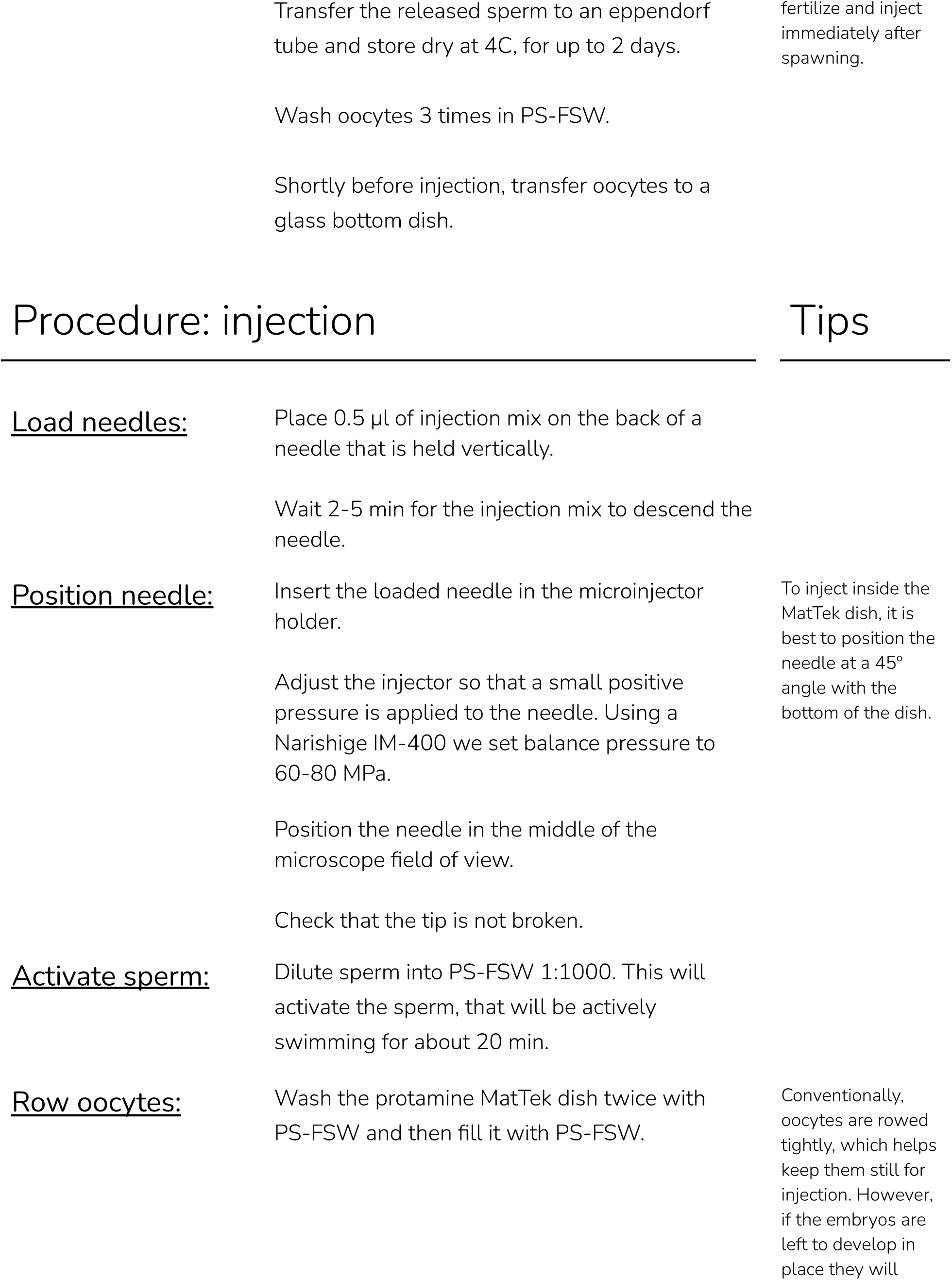
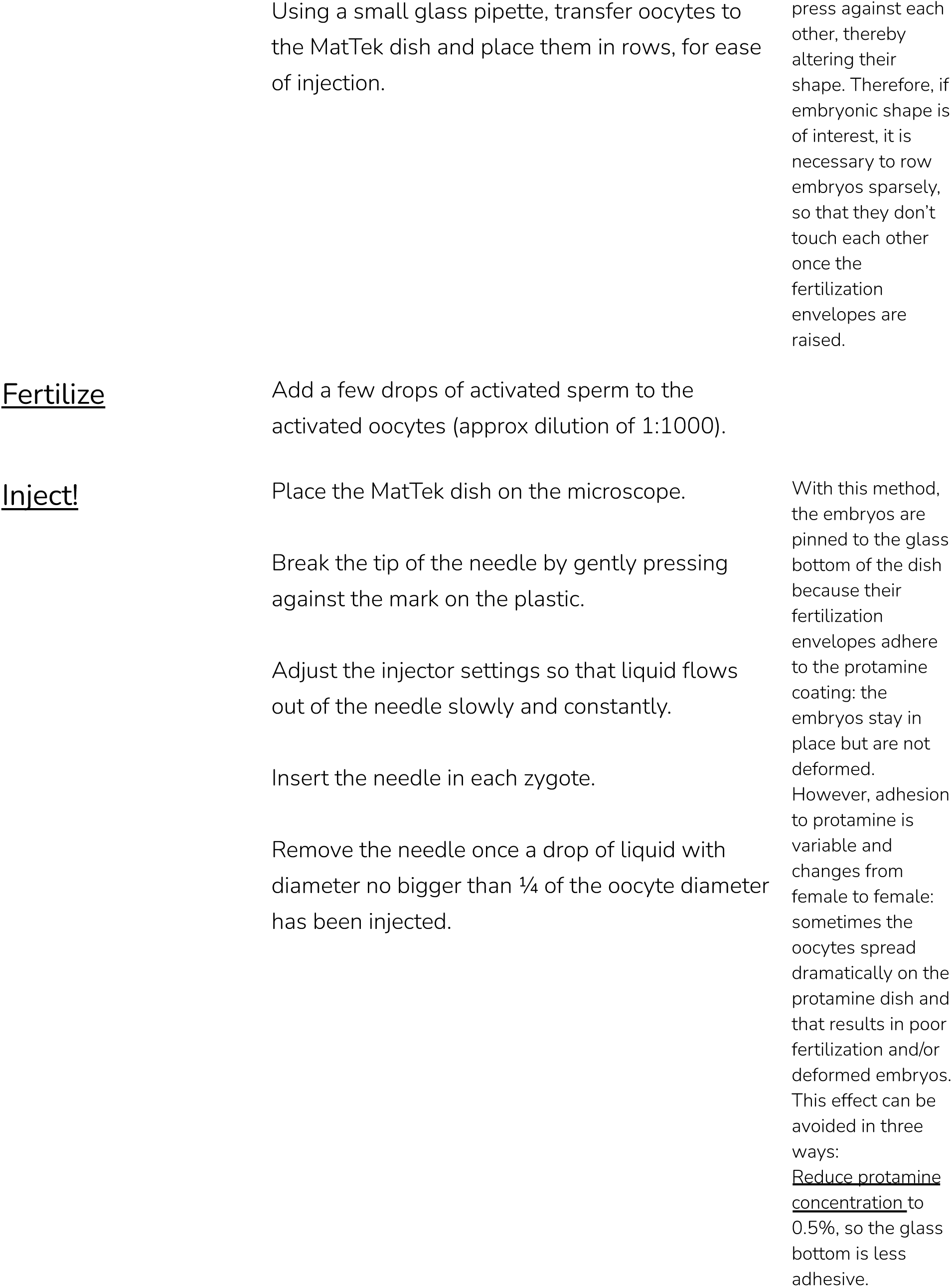
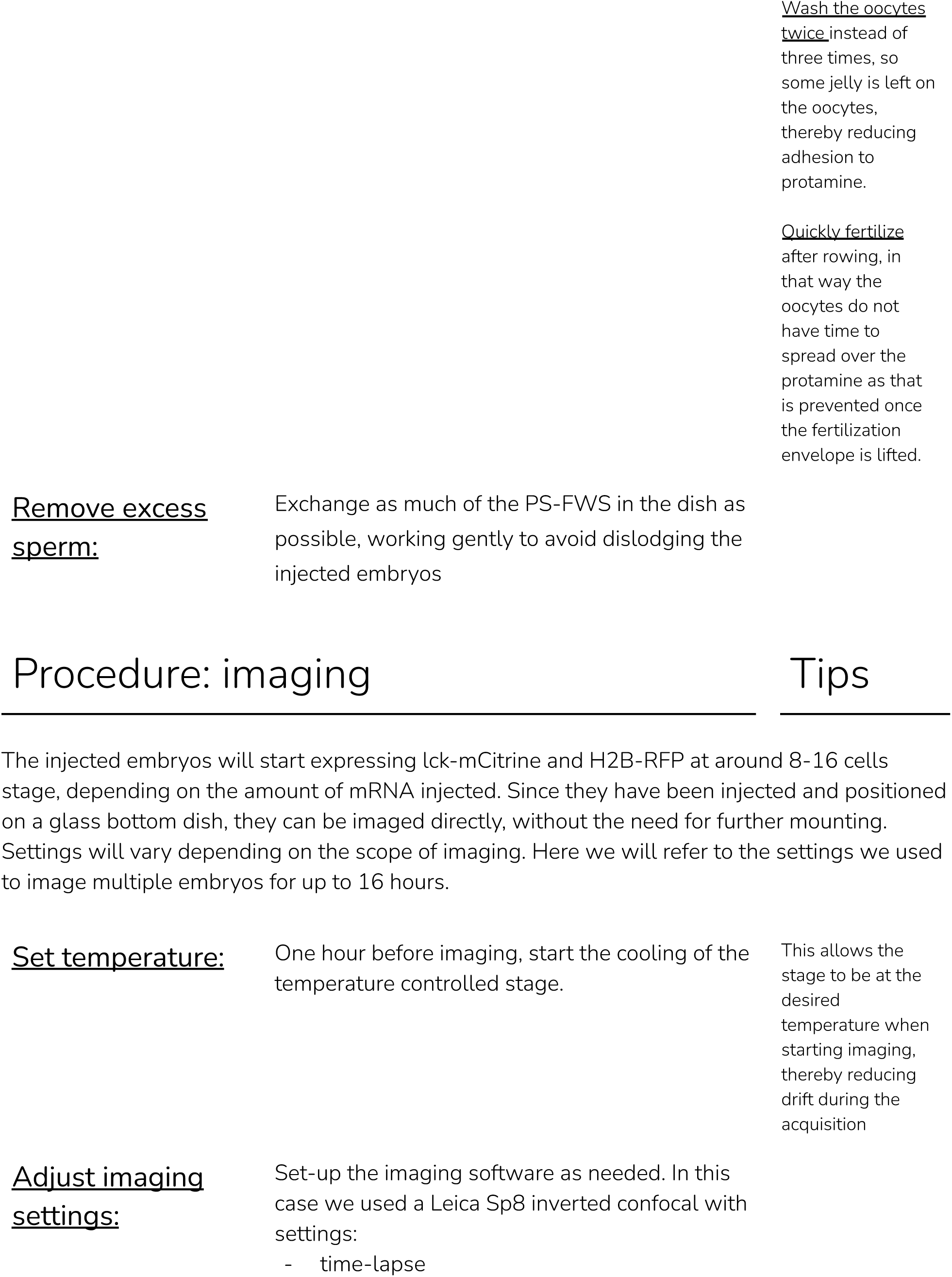
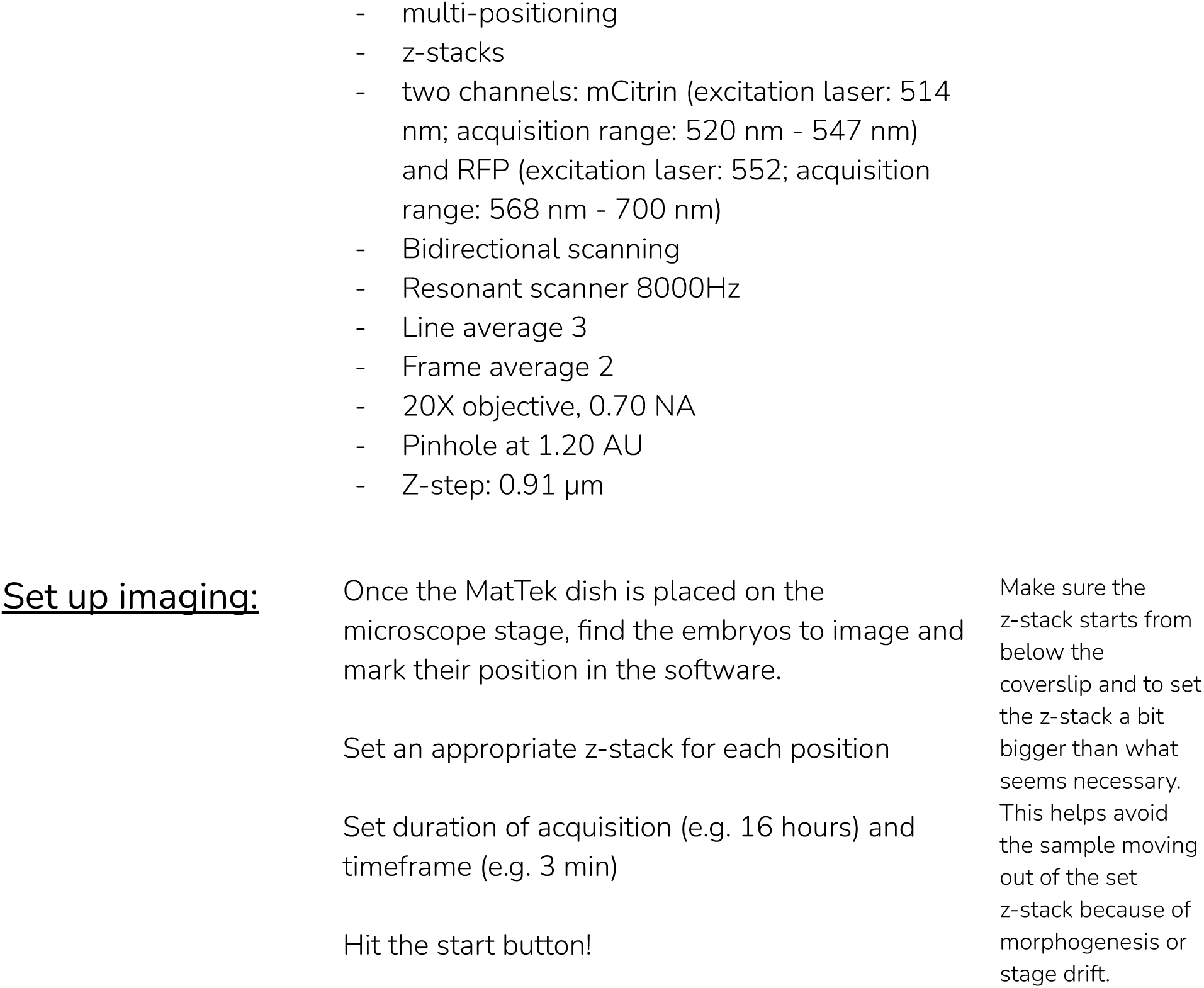

## Results

To analyze how cell shape and cell-cell contacts change during early stages of echinoderm development, we adapted methods for long term live imaging of embryos of three different species, one sea urchin (*L. pictus*) and two asteroid sea stars (*P. miniata* and *P. regularis*). To this aim, we injected 1-cell stage sea urchin embryos with mRNAs coding for a membrane bound mCitrine (lck-mCitrine) and a nuclear CFP (H2B-CFP). Injections were performed directly on glass bottomed petri dishes, so that successfully injected embryos could be imaged directly without further manipulations. Injected embryos were imaged on a confocal microscope starting at the 2-cell stage and until hatching (.ca 10 hours of consecutive imaging; Fig. 5, Movie S3).

Sea star embryos were prepared in a slightly different manner. We injected immature oocytes with mRNA coding for lck-mCitrine and H2B-CFP (or HRAS-GFP and H2B-RFP) and incubated them overnight at 16C. This results in the oocytes translating the mRNAs and expressing the fluorescently tagged proteins (Lénárt et al., 2003; von Dassow et al., 2019). We then activated the oocytes by exposing them to 1-MA for 60 min, which results in the oocytes completing meiosis (Foltz et al., 2004; Jarvela et al., 2014), and fertilized them. Once the embryos reached the 2-cell stage (approximately 3 hpf), we mounted them on a glass bottomed dish (see Methods for details) and imaged them on a confocal microscope. Sea star embryos were imaged from the 2-cell stage and until hatching (.ca 12 hours of consecutive imaging; Fig. 5, Movies S4, S5).

Imaged embryos developed normally as they showed no phenotypes when compared to untreated siblings (Fig. 4O,P). Moreover, the datasets obtained allowed us to perform 3D segmentation of individual cells within the embryos and to track cells over time (Fig. 5). We used the Fiji plug-ins Limeseg (Machado et al., 2019) to segment and track cells. Representative images of the segmentation results are shown in Fig. 5.

## Discussion

The methods described here allow long term live imaging of several echinoderm embryos, and can be easily adapted by most cell and developmental biology labs that wish to expand their work to new model systems. Live imaging of early embryos at subcellular resolution in multiple species will deliver exciting new insight on the evolution of development. A particularly interesting avenue will be that of coupling live imaging with both biophysics and molecular approaches, which has already proven very effective in uncovering mechanisms of morphogenesis in the zebrafish (Behrndt et al., 2012; Maître et al., 2012; Barone et al., 2022, 2017; Krens et al., 2017; Mongera et al., 2018; Munjal et al., 2021), mouse (Maître et al., 2015; Dumortier et al., 2019), Drosophila (Rauzi et al., 2013; Munjal et al., 2015), *C. elegans (Michaux et al*., *2018; Gross et al*., *2019; Chartier et al*., *2021)* and ascidian embryos (Guignard et al., 2020; Godard et al., 2021). Applying these approaches to early development of echinoderms will, for instance, prove invaluable in understanding how a developmental program for epithelialization, i.e. the formation of a blastula, has been modified during evolution, and identifying which cell behaviors (cell shape changes, cell-cell and cell-matrix adhesion, cell division, etc) determine the mode of epithelialization in the different embryos.

Echinoderms also have variation in key aspects of embryonic development at later stages, including gastrulation - with variation in the modes of tissue invagination and cell ingression (Kuraishi and Osanai, 1992; McClay et al., 2020) - and organogenesis, for instance in the organization of neuronal cell types lining the ciliary bands (Hinman and Burke, 2018). The embryos and larvae are still transparent and accessible for imaging at these stages, however the main challenge to live imaging is ciliary movement: as many other marine larvae, echinoderm larvae are excellent swimmers and are propelled by the beating of their cilia, which develop during blastula stages (Gilbert and Barresi, 2017). Live imaging of late blastulae, gastrulae and early larvae, depends on being able to immobilize the specimen without affecting its shape. For short-term imaging (0.5-1h), it is possible to stop ciliary beating by either deciliating - via a quick osmotic shock - or incubating the larvae with drugs inhibiting ciliary movement (Tisler et al., 2016). However, these methods are not suitable to acquire longer timelapses, as they impair normal development (Tisler et al., 2016).

The challenge ahead is therefore to develop mounting and imaging techniques amenable to imaging of echinoderm and other marine embryos at all stages of development. Possible avenues include screening for compounds that inhibit ciliary movement without affecting embryogenesis (Hose, 1985; Semenova et al., 2018), the development of microscopes that can follow the swimming larvae (Krishnamurthy et al., 2020) or of devices that trap the specimen without deforming it (Van Treuren et al., 2019). Overcoming these challenges will open a new chapter for the study of embryonic development, as there is an ocean of morphogenesis waiting to be imaged.

## Supplementary movies legends

**Movie S1 - Timelapse of *Crepidula atrasolea* embryo**. Animal view of C. atrasolea embryo recorded from the 4-cell stage to the 25-cell stage. lck-mCitrine (yellow) and H2B-CFP (blue).

**Movie S2 - Timelapse of *Lytechinus pictus* embryo labeled with vital dye**. Vegetal view of a *L. pictus* embryo recorded from the 1-cell to the 32-cell stage. Membranes were labeled with the vital dye FM4-64.

**Movie S3 - Timelapse of *Lytechinus pictus* embryo labeled by mRNA injection**. Vegetal view of a *L. pictus* embryo recorded from the 32-cell stage to early blastula. lck-mCitrine (green) and H2B-CFP (magenta).

**Movie S4 - Timelapse of *Patiria miniata* embryo labeled by mRNA injection**. Animal view of a *P. miniata* embryo recorded from the 2-cell stage to early blastula. HRAS-GFP (green) and H2B-RFP (magenta). Note the polar bodies are visible in the center of this view.

**Movie S5 - Timelapse of *Patiriella regularis* embryo labeled by mRNA injection**. Lateral view of a *P. regularis* embryo recorded from the 2-cell stage to early blastula. HRAS-GFP (green) and H2B-RFP (magenta).

